## supplementary text 1 for "Network organization of antibody interactions in sequence and structure space: the RADARS model"

Combination of lognormal and exponential distributions (1), with rate of exponential decay λ and variance 2/λ, respectively.

Solution of the integral

$$f(Ksys) = \int_{r=0}^{\infty} \lambda e^{-\lambda r} \frac{\sqrt{\lambda}}{\sqrt{\pi r} 2 \Delta E Ksys} e^{-\frac{\lambda(\Delta G^{\circ} sys)^2}{4 r \Delta E^2}} dr$$

Rearranging, after substituting r=u$^2$

$$f(Ksys) = \frac{2\lambda\sqrt{\lambda}}{\sqrt{\pi}2} \frac{1}{Ksys} \int_{u=0}^{\infty} e^{-\lambda u^2 - \frac{\lambda(\Delta G^{\circ} sys)^2}{4u^2 \Delta E^2}} du$$

Considering ΔE=1 as unit of energy fluctuation

$$f(Ksys) = \frac{2\lambda\sqrt{\lambda}}{\sqrt{\pi}2} \frac{1}{Ksys} \int_{u=0}^{\infty} e^{-\lambda u^2 - \frac{\lambda(\Delta G^{\circ} sys)^2}{4u^2}} du$$

Using integral table identity

$$\int_{z=0}^{\infty} e^{-az^2 - b/z^2} = \frac{1}{2}\sqrt{\frac{\pi}{a}} e^{-2\sqrt{ab}}$$

Substituting $\lambda$ for a and $\lambda/4$ for b

$$\int_{z=0}^{\infty} e^{-\lambda z^2 - \lambda/z^2} = \frac{1}{2}\sqrt{\frac{\pi}{\lambda}} e^{-2\sqrt{\lambda\lambda/4}}$$

Replacing the integral part of the equation

$$f(Ksys) = \frac{2\lambda\sqrt{\lambda}}{\sqrt{\pi}2} \frac{1}{Ksys} \frac{1}{2}\sqrt{\frac{\pi}{\lambda}} e^{-2\sqrt{\frac{\lambda^2(\Delta G^{\circ} sys)^2}{4}}}$$

Simplifying with 2, $\sqrt{\varphi}$, $\sqrt{\pi}$

$$f(Ksys) = \frac{\cancel{2}\lambda\cancel{\sqrt{\lambda}}}{\cancel{\sqrt{\pi}}\cancel{2}} \frac{1}{Ksys} \frac{1}{2}\sqrt{\frac{\cancel{\pi}}{\cancel{\lambda}}} e^{-2\sqrt{\frac{\lambda^2(\Delta G^{\circ} sys)^2}{4}}}$$

gives

$$f(Ksys) = \frac{\lambda}{2} \frac{1}{Ksys} e^{-2\sqrt{\frac{\lambda^2(\Delta G^{\circ})^2}{4}}}$$

Simplifying the exponent

$$f(Ksys) = \frac{\lambda}{2}\frac{1}{Ksys}e^{-\lambda\Delta G°sys}$$

Converting system energy $\Delta G°_{sys}$ to system equilibrium constant Ksys

$$f(Ksys) = \frac{\lambda}{2}Ksys^{-1}e^{-\lambda \ln{(Ksys)}}$$

Simplifying the exponential expression

$$f(Ksys) = \frac{\lambda}{2}Ksys^{-1}Ksys^{-\lambda}$$

$$f(Ksys) = \frac{\lambda}{2}Ksys^{-\lambda-1}$$

Assuming λ=ϕ [golden ratio] and using $\phi+1=\phi^2$

$$f(Ksys) = \frac{\varphi}{2}Ksys^{-\varphi^2}$$

Geometrical interpretation of energy distributions

As antigen is gradually eliminated during the immune response, B cell clones with suboptimal binding to the antigen, e.i. cells growing not exactly in the direction of antigen, will be left without survival signals. Applying this approach in our model a conical network of cells with structurally related antibodies will survive. Because of cross-reactivity with similar structures that share parts of the binding area, antibody secreting cells at a distance from the mean interaction energy surface "cover" a region of shape space, subtending a conical region of interaction space where concentration of the respective antigen is controlled. The covered surface is proportional to the area of a circle with a radius $\Delta G°_{sys}$ corresponding to the distance from $<\Delta G°>_{sys}$, multiplied by a factor $\lambda$ that represents the relationship between conformational diversity and interaction energy in thermodynamic equilibrium.

$$\text{surface area subtended} = \lambda\pi(\Delta G°)^2$$

The relationship of this surface to the total surface of the sphere representing interaction with energy $r\Delta E$ is given by

$$\text{ratio to spherical surface} = \frac{\varphi\pi(\Delta G°)^2}{4\pi r\Delta E^2}$$

The probability of an interaction event, and thereby the survival of a B cell, is given by probability of event = $e^{-\frac{\varphi\pi(\Delta G°)^2}{4\pi r \Delta E^2}}$

Which is a normal distribution, when completed for an integral

$$f(\Delta G°) = \frac{\sqrt{\varphi}}{\sqrt{\pi}2\Delta E} e^{-\frac{\varphi\pi(\Delta G°)^2}{4\pi r \Delta E^2}}$$

Interaction energy is determined by r, the number of bonds between pairs of atoms, and the fluctuation energy of interaction ΔE. We can characterize energy fluctuation as multiples of the energy associated with tonic signaling, since tonic signals apparently arise in the absence of antigen (2) and may therefore represent an intrinsic property of the surface antibody. In the Gaussian function the variance in energy is thus influenced by the number of potential interatomic bonds.

The concentration of antigenic structures available for interaction with given energy is described by the exponential function

f ([Ag])= $\lambda e^{-\lambda \Delta G^2}$

if we assume that immunological mechanisms drive affinity maturation to a point where antigen is optimally eliminated. The rate of this exponential decrease is determined by the rate of probability density change of finding a cell on a given energy shell, which we characterized by $\lambda$. This in turn means that of the energy surfaces determined by the sphere $4\pi r \Delta E^2$ exponentially smaller parts will be actually populated by cells. The integral of the two functions describes the system built from these energy surfaces. The corresponding power law distribution then characterizes the network of interactions.
